# SNMF: Ultrafast, Spatially-Aware Deconvolution for Spatial Transcriptomics

**DOI:** 10.64898/2026.03.17.712043

**Authors:** Luis Alonso, Idoia Ochoa, Ángel Rubio

## Abstract

Sequencing-based spatial transcriptomics has revolutionized the study of tissue architecture, but its ‘spots’ often contain multiple cells, creating a key computational challenge, termed deconvolution, to decipher each spot’s cell-type composition. Reference-free deconvolution methods avoid the need for a matched single-cell RNA-seq dataset, but typically neglect the spatial correlation between neighboring spots and do not leverage modern hardware for efficient computation. Here, we propose SNMF (Spatial Non-negative Matrix Factorization): a rapid, accurate, and reference-free deconvolution method. SNMF extends the standard NMF framework with a spatial mixing matrix that models neighborhood influences, guiding the factorization toward spatially coherent solutions. Our R package is, to our knowledge, the first spatial transcriptomics deconvolution tool to natively support GPU execution, completing benchmark analyses in under one minute—over two orders of magnitude faster than the slowest competing methods— with moderate memory requirements. On synthetic and real benchmark datasets, SNMF significantly outperforms state-of-the-art methods in deconvolution accuracy, and on a human melanoma dataset it recovers biologically meaningful cell-type signatures—including a tumor-boundary transition zone— without any reference input. The proposed mehtod is publicly available at https://github.com/ML4BM-Lab/SNMF.

## Introduction

Next-generation sequencing technologies have transformed gene expression profiling, and spatial transcriptomics has further revolutionized the field by preserving the spatial context of gene expression within intact tissue sections (1), revealing how cellular organization shapes disease progression (2) and function (3) in ways that dissociated single-cell assays cannot.

Sequencing-based platforms (e.g., Visium (4), Slide-seq (5)) can profile thousands of genes without a predefined panel, but each spatial unit—or spot—typically captures transcripts from multiple cells, yielding mixed profiles that confound downstream interpretation. This motivates cell-type deconvolution: inferring the cell-type composition of each spot *in silico* (6–8). Reference-free methods avoid the need for matched scRNA-seq datasets (9), but most ignore the spatial structure of the data entirely—despite the well-established influence of neighboring spots on expression profiles (7). Among state-of-the-art reference-free methods, only a minority leverage spatial coordinates (Table 1) (10). A further limitation is computational: no R-based deconvolution method natively exploits GPU acceleration, despite the scale of modern datasets (11).

**Table 1.** Comparison of state-of-the-art deconvolution methods for spatial transcriptomics that can operate without a single-cell reference, ordered by citation count as of February 2026. Methods are characterized by their programming language, use of marker genes (MGs) for deconvolution (not annotation), whether they produce annotated output, whether they recover all cell types (CTs) present, package availability, GPU support, and use of spatial coordinates. SNMF is the only R-based method that combines native GPU support with explicit spatial modeling.

| Method | Language | Uses MGs | Annotates | All CTs | Package | Uses GPU | Spatial coords. |
| --- | --- | --- | --- | --- | --- | --- | --- |
| CARD<sup>1</sup> (2022, *Nat. Biotechnol.*) (7) | R | Yes | No | — | Yes | No | Yes |
| STdeconvolve (2022, *Nat. Commun.*) (6) | R | No | Yes | No | Yes | No | No |
| SpiceMix (2023, *Nat. Genet.*) (21) | Python | No | No | — | Yes | Yes | Yes |
| Starfysh<sup>2</sup> (2025, *Nat. Biotechnol.*) (22) | Python | No | Yes | Yes | Yes | Yes | No |
| BayesTME (2023, *Cell Syst.*) (23) | Python | No | No | — | Yes | No | Yes |
| SMART (2024, *Genome Biol.*) (24) | R | Yes | Yes | Yes | Yes | No | No |
| RETROFIT (2023, *bioRxiv*) (25) | R | No | Yes | Yes | Yes | No | No |
| *Not included in benchmark:* |  |  |  |  |  |  |  |
| UniCell Deconvolve<sup>3</sup> (2023, *Nat. Commun.*) (26) | Python | No | Yes | No | Yes | Yes | No |
| MUSTANG<sup>4</sup> (2024, *Cell*) (27) | Python/R<sup>5</sup> | No | No | — | No | No | Yes |
| CellsFromSpace<sup>4</sup> (2024, *Bioinformatics*) (28) | R | No | No<sup>6</sup> | — | No | No | No |
| SNMF (this work) | R | No | No | — | Yes | Yes | Yes |

<sup>1</sup> CARD is primarily a reference-based method that requires a matched scRNA-seq dataset; here we use its reference-free mode, which accepts marker genes instead of a single-cell reference.
<sup>2</sup> Starfysh accepts optional marker genes; if not provided, it derives cell-type signatures via archetypal analysis. In our benchmark, Starfysh was run without user-provided marker genes.
<sup>3</sup> Excluded: zero-shot method with a fixed set of predefined cell types; the user cannot specify $k$ .
<sup>4</sup> Excluded: no usable software package available at the time of benchmarking.
<sup>5</sup> MUSTANG uses both R and Python for different pipeline steps and does not support running the full pipeline in either language alone.
<sup>6</sup> CellsFromSpace does not automatically annotate independent components but provides an interactive app for guided manual annotation.

To address these three limitations simultaneously, we propose **SNMF** (Spatial Non-negative Matrix Factorization): a rapid, accurate, and reference-free deconvolution method that extends standard NMF with a spatial mixing matrix **S** modeling neighborhood influences. SNMF is, to our knowledge, the first spatial transcriptomics deconvolution tool in the R ecosystem with native GPU support, completing benchmark analyses in under one minute. On synthetic and real benchmark datasets, it significantly outperforms state-of-the-art methods, and on a human melanoma dataset it recovers biologically meaningful signatures—–including a tumor-boundary transition zone—–without any reference input.

## Methods

### A. SNMF algorithm

SNMF deconvolves a spatial transcriptomics count matrix by extending the standard NMF framework with a spatial mixing matrix. Let 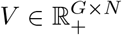 denote the observed gene expression matrix, where *G* is the number of genes and *N* the number of spots. Standard NMF approximates *V* ≈ *WH*, where 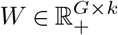 contains the expression profiles of *k* cell types and 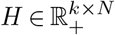 encodes cell-type proportions per spot. SNMF augments this with a fixed spatial mixing matrix 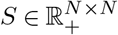:

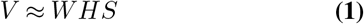

Tissue organization follows biological patterns where cells of the same type tend to cluster together (12–14), and spatial autocorrelation of marker genes has been reported (7). The matrix *S* encodes this neighborhood structure via a Gaussian kernel: 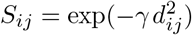, where *d*_*ij*_ is the Euclidean distance between spots *i* and *j*. Entries below 10^−3^ are set to zero, and each row is *f*_1_-normalized. Rather than treating *γ* as a free parameter, we set it so that the mean diagonal of the normalized *S* equals a target *τ* (default *τ* = 0.5), meaning each spot contributes equally with its local neighborhood on average. The optimal *γ* is found by BFGS minimization (15–18). *S* is computed once before optimization and remains fixed throughout.

Because spatial transcriptomics data are count-based, SNMF minimizes the KL divergence between *V* and *WHS* (full derivation in Supplementary Section 1):

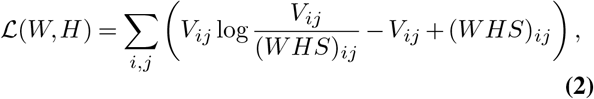

which is equivalent to fitting a Poisson model with mean (*WHS*)_*ij*_ (19). While sequencing data often exhibit overdispersion, the KL objective remains robust in practice and avoids the sensitivity of Euclidean loss to high-count genes. An ablation study minimizing the Frobenius norm instead resulted in performance degradation (see Supplementary Figure 5). Following Lee & Seung (19), we minimize ℒ using multiplicative update rules (derivation in Supplementary Section 2):

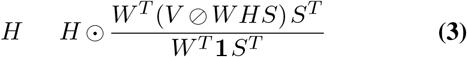

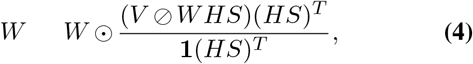

where ⊙ and ⊘ denote element-wise multiplication and division, and **1** is a matrix of ones. The *W* update is identical to standard KL-NMF with *H* replaced by *HS*. Both updates consist entirely of matrix multiplications and element-wise operations, making them directly amenable to GPU acceleration.

To resolve the scaling ambiguity inherent in NMF (*WD* · *D*^−1^*H* is equally valid for any positive diagonal *D*), columns of *W* are rescaled to have identical 75th percentiles, with *H* rescaled by the inverse factors. Rows of *HS* are then normalized to sum to one, yielding relative cell-type proportions per spot. *W* and *H* are initialized from a uniform distribution; 10 independent restarts are run, and after a burn-in of one-tenth of the total iterations, the best restart (lowest ℒ) continues to convergence (max 2,000 iterations or relative change *<* 10^−4^). The number of cell types *k* is specified by the user. SNMF is implemented in R using GPUmatrix (20), with seamless CPU fallback. Although *S* is sparse after thresholding, it is stored as a dense matrix for faster GPU operations; for very large *N* (e.g., *>* 50,000 spots), sparse representations would be needed.

### B. Datasets

We benchmarked SNMF on four spatial transcriptomics datasets spanning synthetic and real data, with varying numbers of cell types and tissue contexts.

#### Pancreatic ductal adenocarcinoma (PDAC)

We generated synthetic spatial transcriptomics data using scDesign3 (29), a simulator that models gene expression across spatial coordinates while respecting user-specified cell-type proportions. As a reference, we used a publicly available PDAC scRNA-seq dataset (30) with 20 annotated cell types. We retained the 2,000 most highly variable genes. The resulting dataset comprises 428 spots arranged on a regular grid, with known ground-truth cell-type proportions at every location, enabling direct evaluation.

#### Triple-negative breast cancer (TNBC)

We used the synthetic dataset generated by He *et al*. (22), based on a TNBC scRNA-seq reference (31). The generation process mimics a dense, square ST-like grid composed of 2,500 spots. Five cell types were simulated with spatial proportion maps drawn from independent two-dimensional Gaussian processes, yielding smooth yet heterogeneous spatial patterns. Ground-truth proportions are available at every spot.

#### Dorsolateral prefrontal cortex (DLPFC)

We used 12 human postmortem DLPFC tissue sections (32) profiled by 10x Visium (4), each manually annotated into cortical layers and white matter (WM) by the original authors. Eight sections (151507–151510, 151673–151676) contain annotations for all six cortical layers plus WM (*k* = 7); the remaining four sections (151669–151672) lack Layers 1 and 2 annotations and were analyzed with *k* = 5. Genes expressed in fewer than 10% of spots within a section were removed; the number of spots and genes retained per section are reported in Supplementary Table 4. This dataset is the *de facto* benchmark for spatial domain identification methods and provides a stringent test on real data.

#### Melanoma sample

We applied SNMF to a spatial transcriptomics dataset of a lymph node metastasis sample (33) from a patient diagnosed with stage III melanoma, for biological validation. This sample was sequenced using the ST technology (34), and is accompanied by histological annotations on a hematoxylin and eosin-stained (H&E) image from one of the two consecutive sections used for the spatial arrays.

This annotation was performed manually by a trained pathologist, who identified melanoma, stromal and lymphoid tissue. Signatures to annotate spots belonging to these regions were published by Zhao et al. (35) and are listed in Supplementary Table 6. A characterization of a transcriptionally distinct tumor-boundary transition zone is available from the original study, providing an independent reference for assessing the biological plausibility of the inferred **W** and **H** matrices. As no continuous ground-truth proportions are available, this dataset is used for qualitative validation only and is not included in the quantitative benchmarking.

### C. Benchmarking framework

We compared SNMF against seven methods that operate without a scRNA-seq reference (Table 1): STdeconvolve (6), SpiceMix (21), Starfysh (22), BayesTME (23), SMART (24), RETROFIT (25), and CARD (7) (reference-free mode with marker genes). Methods requiring marker genes received lists derived using Seurat’s `FindAllMarkers` (36) (logFC ≥ 0.25, min.pct = 0.25, Wilcoxon test, adjusted *p <* 0.05); the same lists were used for all methods per dataset (Supplementary Tables **??**).

On synthetic datasets, predicted components were matched to ground-truth cell types via the Hungarian algorithm (37) on Pearson correlations, and accuracy was measured by Root Mean Squared Error (RMSE). On DLPFC, performance was assessed by ARI between dominant cell type assignments and expert annotations; this evaluates spatial domain recovery rather than deconvolution accuracy *per se*. All methods used *k* equal to the number of annotated regions. Execution time and memory were benchmarked on a GeForce RTX 3090 Ti GPU (24 GB VRAM).

## Results

### D. Spatial regularization stabilizes factorization in high-dimensional settings

A central contribution of SNMF is the spatial mixing matrix **S**, which encodes neighborhood influences o n e ach s pot’s e xpression p rofile. To quantify its effect, we performed an ablation study varying the target mean diagonal *τ* from 0.1 (strong spatial smoothing) to 1.0 (no smoothing; **S** = **I**, equivalent to standard NMF). Full results are reported in Supplementary Fig. 4.

The effect of spatial regularization was strongly dependent on the number of cell types. For the TNBC dataset (*k* = 5), performance was largely stable across all values of *τ*, with the identity configuration (**S** = **I**) performing comparably to configurations with active smoothing (Supplementary Fig. 4A). This indicates that for small *k*, the 10 random restarts employed by SNMF are sufficient to find a good factorization without spatial guidance, and that the spatial term neither helps nor hurts in this regime.

For the PDAC dataset (*k* = 20), configurations with *τ <* 1.0 consistently outperformed the identity, with the default value of *τ* = 0.5 yielding the best or near-best RMSE (Supplementary Fig. 4B). This demonstrates that **S** acts as a regularizer that guides the optimization toward better solutions in high-dimensional factorization landscapes, where the search space is large and random initialization alone is less reliable.

### E. SNMF accurately deconvolves spatial transcriptomics data

On the TNBC dataset (*k* = 5 cell types), SNMF achieved a median RMSE of 0.055 (IQR: 0.034–0.069), compared to 0.081 for the next best-performing method, STdeconvolve (*p <* 0.001, Wilcoxon signed-rank test, Bonferroni-corrected). The improvement was statistically significant against all seven competitors, and SNMF displayed a consistently tighter interquartile range, indicating more robust accuracy across individual spots (Fig. 2B).

**Fig. 1.**
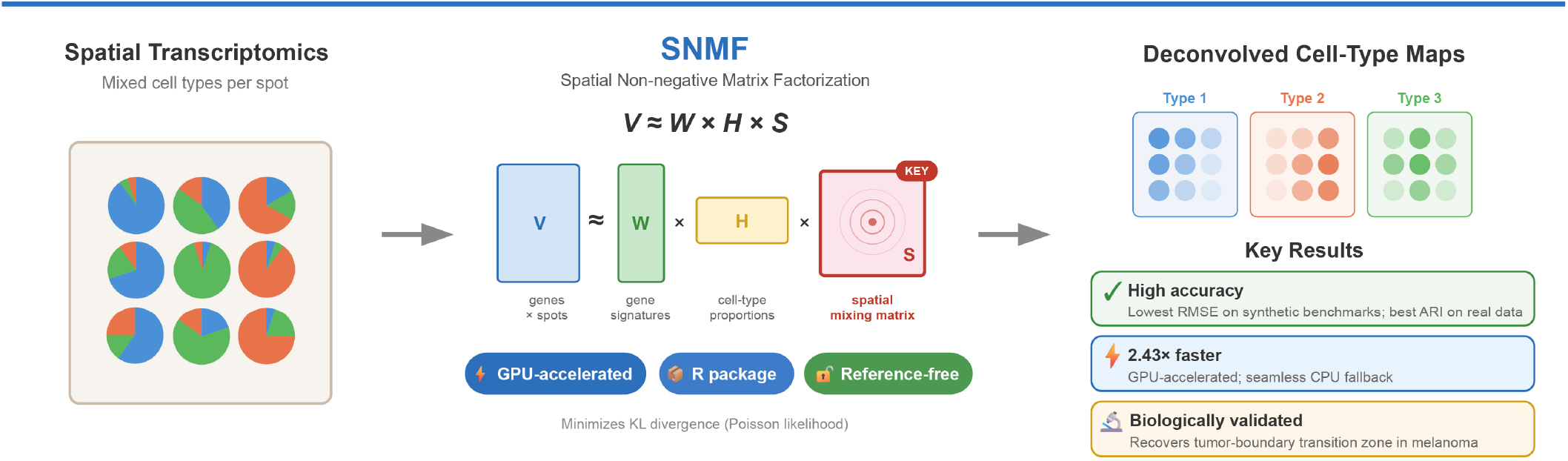
Overview of SNMF for spatial transcriptomics deconvolution. Spatial transcriptomics spots contain mixtures of cell types. SNMF decomposes the gene expression matrix *V* into gene signatures *W*, cell-type proportions *H*, and a spatial mixing matrix *S*, enabling recovery of spatially resolved cell-type maps.

**Fig. 2.**
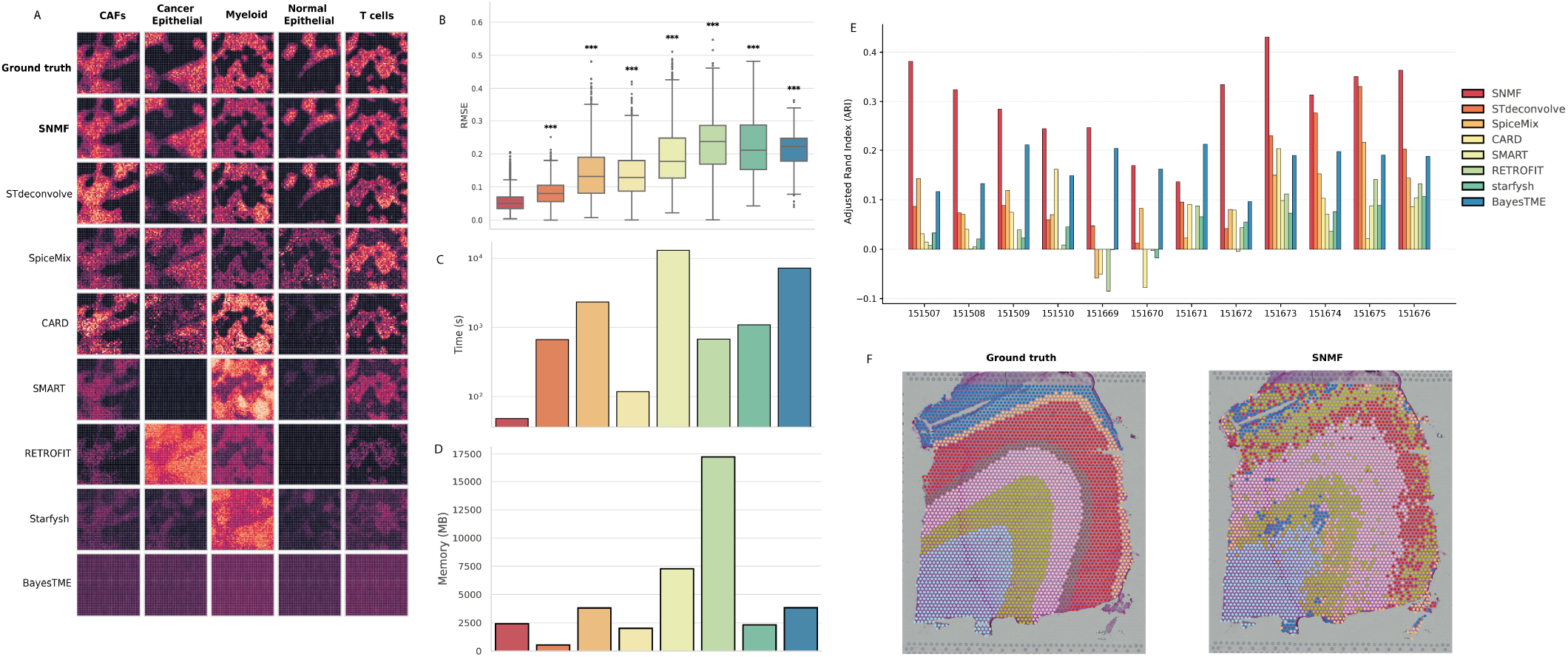
Benchmarking SNMF against state-of-the-art deconvolution methods. (**A)** Spatial heatmaps of predicted cell-type proportion maps for all benchmarked methods and the ground truth on the TNBC dataset. Each row corresponds to a cell type and each column to a method; color intensity represents predicted proportion on the same scale within each row. SNMF most faithfully recapitulates the ground-truth spatial patterns, while competing methods show increased noise, missed cell populations, or blurred boundaries. **(B)** Deconvolution accuracy on the TNBC dataset measured by Root Mean Square Error (RMSE) between predicted and ground-truth cell-type proportions across all spots. SNMF achieves a significantly lower median RMSE and tighter interquartile range than all competing methods. Significance levels relative to SNMF are indicated above each box (ns: *p* ≥ 0.05; *: *p <* 0.05; **: *p <* 0.01; ***: *p <* 0.001; Wilcoxon signed-rank test, Bonferroni-corrected). **(C)** Computational runtime (seconds, log scale) on the TNBC dataset. SNMF (GPU) completes in 48s, a 2.43-fold speedup over the next fastest method. **(D)** Peak RAM usage (MB) for each method on the same hardware as (C). SNMF requires 2.36 GBs, substantially less than memory-intensive methods such as RETROFIT (16.83 GBs). **(E)** ARI across the 12 DLPFC tissue sections for each method. SNMF achieves the highest mean ARI of 0.298 *±* 0.083. **(F)** Spatial maps of the predicted (top) and expert-annotated (bottom) dominant cel -type for a representative DLPFC section (sample 151674). Each spot is colored by its assigned dominant cell type. SNMF’s predicted spatial organization closely matches the expert annotation, with cortical layers and white matter clearly delineated.

The spatial quality of the deconvolution is illustrated in Fig. 2A. SNMF’s predicted cell-type proportion maps closely recapitulate the ground-truth spatial patterns for all five cell types, preserving both the sharp boundaries between regions and the relative intensity differences across the tissue. Among competing methods, STdeconvolve and SpiceMix recover the broad spatial organization but produce noisier maps, particularly for the T cell population. CARD appears to conflate Cancer Epithelial signal with other cell types, yielding an overestimated Cancer Epithelial component and weak recovery of Normal Epithelial and T cells. SMART fails to resolve four of the five cell types, producing near-blank maps for all but one component. RETROFIT produces spatially noisy maps across all cell types, with speckled patterns that do not match the ground-truth structure. Starfysh yields diffuse, low-contrast maps that lack the spatial specificity of the ground truth. BayesTME produces uniform maps with no discernible spatial structure, suggesting convergence failure on this dataset.

Similar results were obtained on the PDAC dataset (*k* = 20 cell types). SNMF achieved the lowest median RMSE across all methods (Supplementary Fig. 6A), with the performance gap widening relative to the TNBC results—consistent with the ablation study (Supplementary Fig. 4), which shows that spatial regularization is most beneficial in high-dimensional settings. The full inferred spatial maps are shown in Supplementary Fig. 7–8.

This section also illustrates the advantage of using the KL divergence instead of the Frobenius norm: the RMSE in the predictions of the proportions for each cell type is significantly smaller (Wilcoxon paired-test p.value < 10^−^16) for both datasets (Supplementary Fig. 5).

### F. GPU acceleration substantially reduces computation time

We benchmarked execution time and peak memory usage on the TNBC and PDAC datasets. SNMF with GPU acceleration completed in 48 s on the TNBC dataset, representing a 2.43-fold speedup over the next fastest method (CARD, 117 s) and two to three orders of magnitude faster than the slowest methods (SMART, 13,007 s; BayesTME, 7,180 s) (Fig. 2C). Note that the time axis is on a logarithmic scale, so the visual gap between SNMF and the slowest methods substantially understates the absolute runtime difference. The majority of competing methods required between 10 and 80 minutes, placing SNMF’s runtime one to two orders of magnitude below the field average. Runtime results on the PDAC dataset show a similar pattern (Supplementary Fig. 6B).

Peak memory usage is shown in Fig. 2D for TNBC. SNMF required approximately 2.36 GB, placing it in the lower half of the methods tested. STdeconvolve exhibited the smallest memory footprint (∼ 0.53 GB), while RETROFIT required ∼16.83 GB—an order of magnitude more than most methods and potentially prohibitive on standard hardware. SNMF’s memory usage is dominated by the dense *N × N* spatial mixing matrix **S** and the GPU-resident copies of **W, H**, and intermediate products; for the dataset sizes tested here, this remains well within the capacity of consumer-grade GPUs (typically 8–24 GB VRAM). Similar results were obtained for PDAC (Supplementary Fig. 6C).

### G. SNMF demonstrates spatial domain recovery on real data

We evaluated the considered methods on the human DLPFC tissue sections (32) using the ARI metric, as continuous ground-truth proportions are unavailable. Mean ARI values across the 12 sections are shown in Fig. 2E, with full per-sample numerical results reported in Supplementary Table 5. SNMF achieved the highest ARI in 11 out of 12 samples, with a mean ARI of 0.298 *±* 0.083 (SD) across all sections. The next best-performing methods were BayesTME (mean ARI: 0.171 *±* 0.037) and STdeconvolve (mean ARI: 0.129 *±* 0.099). The difference between SNMF and each competitor was statistically significant (*p <* 0.001, paired Wilcoxon signed-rank test across the 12 sections). In the section where SNMF did not rank first—sample 151671— it ranked second; these section belongs to the group lacking Layer 1 annotation (*k* = 5), which may reduce discriminative power.

We note that absolute ARI values are moderate for all methods (Supplementary Table 5), reflecting the inherent difficulty of this evaluation setting: each annotated layer contains a mixture of cell types, and the majority-voting strategy penalizes methods that correctly predict heterogeneous compositions within a layer. The ARI should therefore be interpreted as a relative ranking of methods rather than an absolute measure of deconvolution quality. Nevertheless, the spatial maps of annotated vs predicted dominant cell types demonstrate that SNMF achieves better spatial domain recovery than other methods (see Supplementary Fig. 9 for a full comparison and Fig. 2F for a representative section with SNMF).

### H. SNMF recovers biologically meaningful cell-type signatures

To assess whether SNMF’s inferred matrices capture genuine biological signal, we applied it on the human melanoma dataset with *k* = 4. Even though the manual annotations distinguish three regions (stroma, lymphoid and melanoma tissue), they do not cover the whole slide, leaving some regions unlabeled (Fig. 3A). Moreover, Zhao et al. (35) provided signatures for four cell-types. Hence, *k* = 4 was the natural selection.

**Fig. 3.**
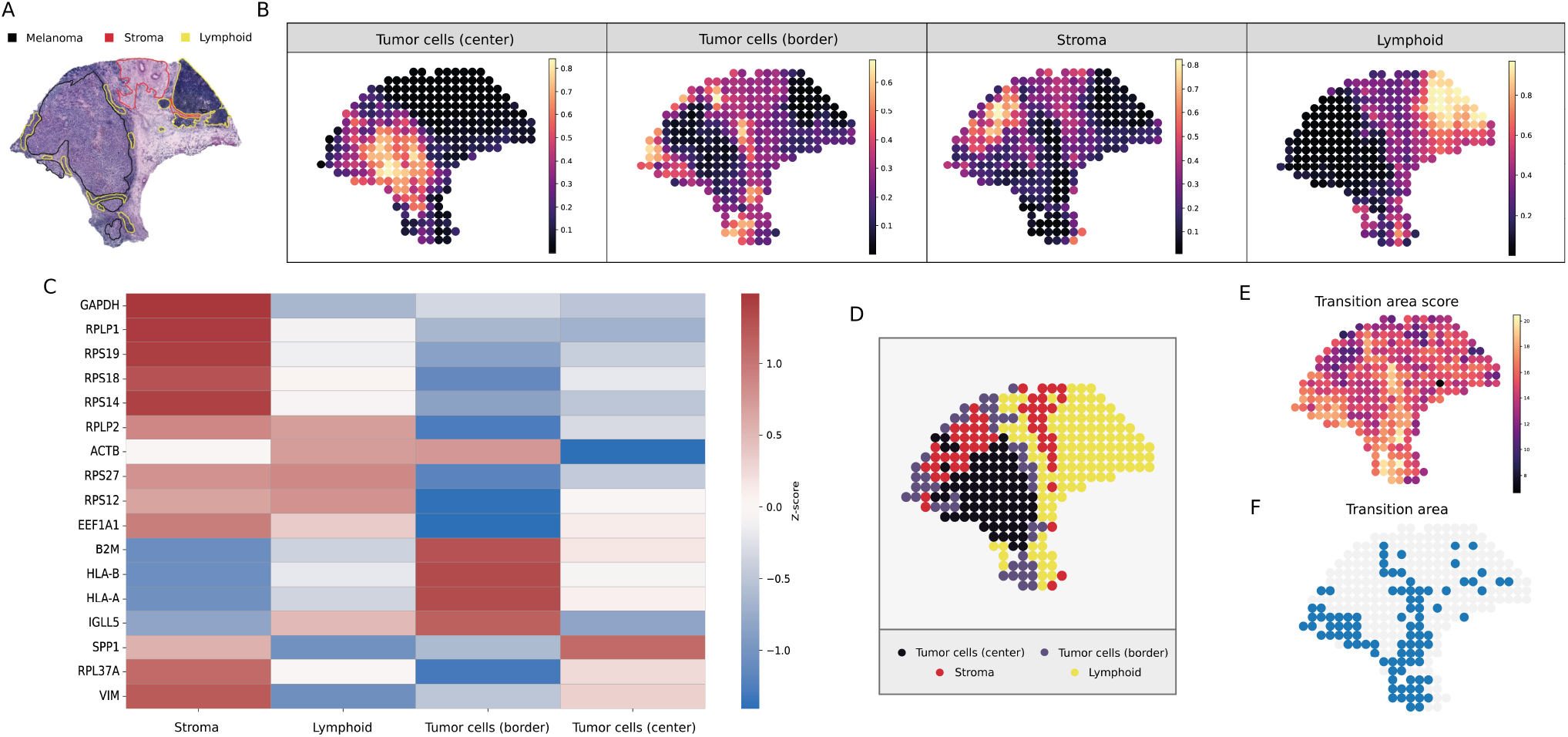
Biological validation of SNMF on a human melanoma spatial transcriptomics dataset (33). **(A)** Histological annotations of melanoma, stroma and lymphoid tissue. **(B)** Spatial maps of the inferred cell-type abundance matrix **H** across the melanoma tissue section. Each panel shows the predicted proportion of one annotated cell type. **(C)** Heatmap of the gene loadings (columns of **W**) for each component, restricted to the top 4 genes with the highest loadings. **(D)** Dominant SNMF component mapping for each spot. **(E)** “Transition-area” score obtained by summing up the log-normalized counts of every gene in the signature from the original study. **(F)** Transition region defined by thresholding the “transition-zone” score so that spots with more than 16 total log-normalized counts are set to be part of this region.

To annotate SNMF components (matrix **W**), we computed enrichment scores for the available cell-type signatures marker gene sets. For each component, gene loadings were standardized, and the mean z-score of marker genes was compared to the background distribution of all genes. Cell types were assigned to the component with the highest enrichment score exceeding a predefined threshold, enforcing a one-to-one mapping between components and cell types. This assignment resulted in no ambiguous components.

The annotated cell types showed strong spatial concordance with the histologically defined regions (Fig. 3B). In particular, components annotated as tumor cells (center) and tumor cells (border) recapitulated the spatial organization of the tumor mass, with central tumor signatures concentrated in the tumor core and border tumor signatures enriched at the tumor–stroma interface. Likewise, stromal and lymphoid components localized to regions consistent with their histological annotations, indicating that the SNMF decomposition captures biologically meaningful spatial patterns.

Inspection of the genes driving each component further supported this interpretation (Fig. 3C). Ribosomal and house-keeping genes (e.g., *RPLP1, RPS19, EEF1A1*) were enriched in stromal and lymphoid components, while antigen presentation genes such as *HLA-A* and *HLA-B* were strongly associated with the tumor-border component, consistent with the immune activity typically observed at tumor–stroma interfaces.

Mapping the dominant SNMF component for each spot revealed a clear spatial partitioning of the tissue (Fig. 3D). Tumor-center spots formed a contiguous region corresponding to the melanoma core, while tumor-border spots delineated the interface between tumor and surrounding tissue. Stromal and lymphoid components occupied adjacent but distinct regions, reflecting the heterogeneous microenvironment surrounding the tumor.

Leveraging the transition-zone signature, we defined a “transition-zone” score by summing up the expression of all genes in the signature per spot (Fig. 3E). Applying a threshold of 16 log-normalized counts, we could categorically define this transition zone (Fig. 3F). The transition score in this transition area was positively correlated with the proportions of border tumor cells (*ρ* = 0.5624, *p <* 0.005), supporting the reliability of the SNMF inferred proportions. Given that transition areas generally tend to be heterogeneous and that this one has been clustered as an independent component in the original study, we tested whether this transition score was also correlated with the proportions entropy, yielding a positive correlation (*ρ* = 0.2018, *p* = 0.05).

These results demonstrate that SNMF recovers interpretable and biologically grounded transcriptional signatures without any reference data or marker gene input, and that the inferred **W** matrix can be used directly for downstream cell-type annotation.

## Conclusion

We have presented SNMF, a reference-free deconvolution method for sequencing-based spatial transcriptomics. Its central contribution is the spatial mixing matrix **S**, which encodes neighborhood influences directly within the NMF objective at negligible computational cost. Our ablation study shows that **S** acts as a regularizer that is most beneficial when the number of cell types is large—precisely the setting where deconvolution is most challenging—while incurring no accuracy penalty when it is not needed, making *τ* = 0.5 a robust default. Implemented as an R package with native GPU support, SNMF completes benchmark analyses in under one minute and, on a human melanoma dataset, recovers biologically interpretable cell-type signatures—including a tumor-boundary transition zone—without any reference input.

Several limitations should be noted. The dense *N × N* storage of **S** limits scalability to datasets with tens of thousands of spots; sparse representations are a natural extension. The Poisson likelihood implied by the KL divergence does not account for overdispersion, though the multiplicative update framework generalizes to a negative binomial likelihood with gene-specific dispersion parameters, which may improve recovery of rare cell types. The spatial term does not improve accuracy for small *k*, and the method was validated on a limited number of datasets. Future work includes learning the bandwidth parameter end-to-end, extending to three-dimensional spatial data, and optionally incorporating single-cell references when available.

## Data availability

All the code and scripts used in this study are available at https://github.com/ML4BM-Lab/SNMF-paper/ and accompanying data in https://doi.org/10.5281/zenodo.18852117.

## Supplementary Note 1: Derivation of the Kullback-Leibler Divergence

We derive the closed-form expression of the Kullback–Leibler (KL) divergence between two Poisson distributions *p* and *q* with parameters *λ*_1_ and *λ*_2_, respectively.

Let *X* be a discrete random variable. A Poisson distribution with parameter *λ >* 0 is defined by the probability mass function

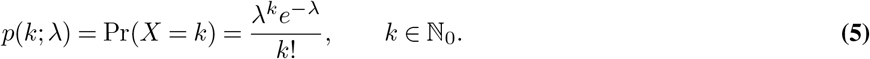

The Kullback–Leibler divergence between two discrete distributions *p* and *q* defined on the same support is

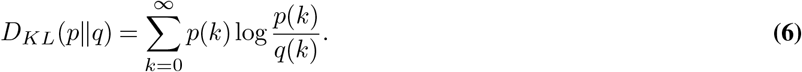

For two Poisson distributions *p*(*k*; *λ*_1_) and *q*(*k*; *λ*_2_), the ratio between their probability mass functions is

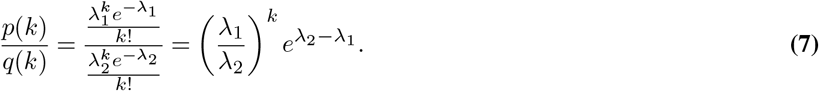

Taking the logarithm gives

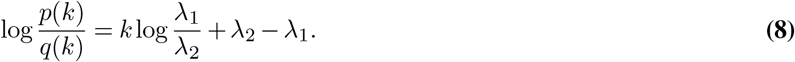

Substituting this expression into the definition of the KL divergence yields

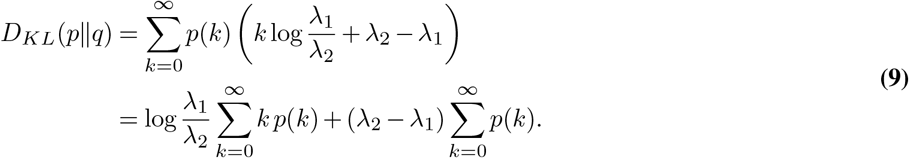

For a Poisson distribution, the expectation satisfies

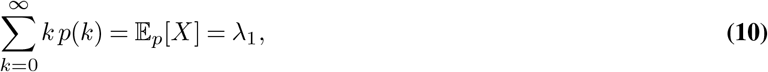

and by normalization

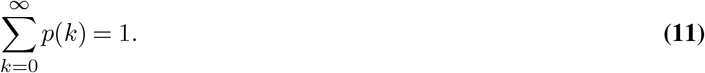

Therefore, the KL divergence between two Poisson distributions is

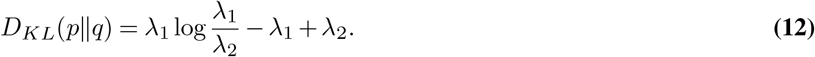

If we substitute the parameters *λ*_1_ and *λ*_2_ by the observed and predicted entries of the matrices *V* and *WHS*, respectively, we obtain the element-wise KL divergence used in SNMF:

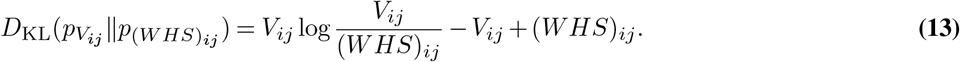

## Supplementary Note 2: SNMF: Derivation of Multiplicative Update Rules

We derive the multiplicative update rules for the abundance matrix *H* and the basis matrix *W* by minimizing the KL divergence between the observed count matrix *V* and its approximation *WHS*. We follow the framework of Lee & Seung (19), which splits the gradient of the objective into positive and negative parts to construct updates that guarantee non-negativity.

### Objective function

The objective is to minimize:

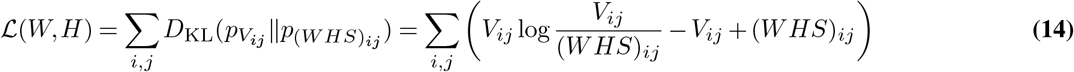

where *i* indexes genes (*i* = 1,…, *G*) and *j* indexes spots (*j* = 1,…, *N*). The matrix 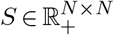 is fixed throughout optimization. We seek non-negative matrices 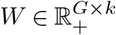 and 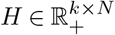 that minimize ℒ.

The general multiplicative update rule for a parameter *θ* is obtained by decomposing the gradient ∇_*θ*_ℒ into its positive and negative parts, 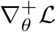 and 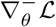, such that 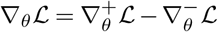 with both parts non-negative, and applying:

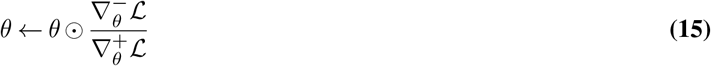

This update sets ∇_*θ*_ℒ = 0 at a fixed point and preserves non-negativity provided *θ* is initialized with non-negative values.

### Update rule for *H*

We compute the partial derivative of ℒ with respect to an element *H*_*ab*_:

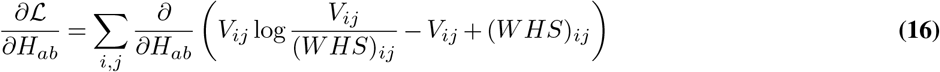

Since *V*_*ij*_ does not depend on *H*_*ab*_, only the last two terms contribute:

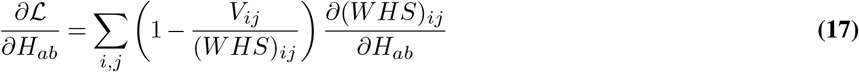

The derivative of the approximation term with respect to *H*_*ab*_ is:

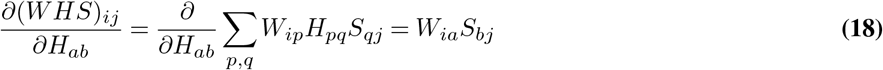

since only the term with *p* = *a* and *q* = *b* survives. Substituting:

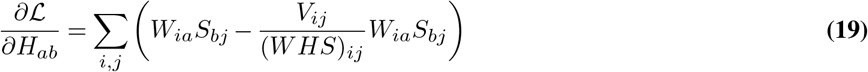

Separating into positive and negative parts and writing in matrix form:

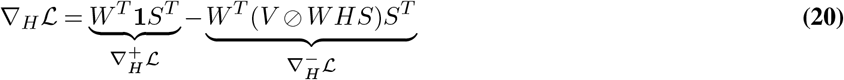

where **1** is a *G × N* matrix of ones and ⊘ denotes element-wise division. Both parts are non-negative since *W, S, V*, and *WHS* are all non-negative. Applying the multiplicative update framework:

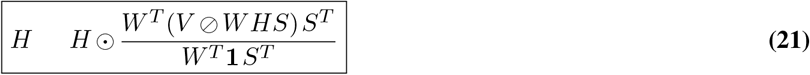

### Update rule for *W*

We compute the partial derivative of ℒ with respect to an element *W*_*ab*_:

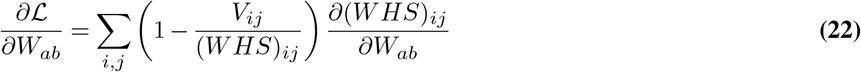

The derivative of the approximation term is:

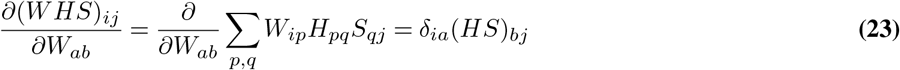

where *δ*_*ia*_ is the Kronecker delta, which restricts the sum to rows *i* = *a*. Substituting and summing over *j* only (since *δ*_*ia*_ eliminates the sum over *i*):

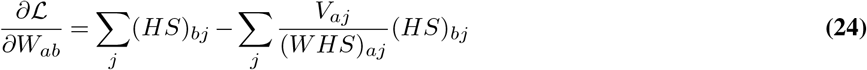

Writing in matrix form and separating into positive and negative parts:

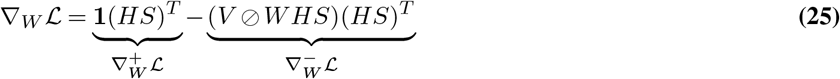

Applying the multiplicative update framework:

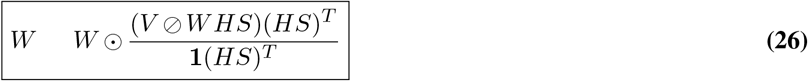

This result is identical to the Lee & Seung update rule for standard NMF under KL divergence, with *H* replaced by *HS* throughout — a direct consequence of the linear role of *S* in the model.

### Convergence and non-negativity

Both updates preserve non-negativity: if *W* and *H* are initialized with strictly positive values, the multiplicative structure ensures they remain strictly positive at every iteration. The denominators are always positive under the same condition.

We now argue that the updates are non-increasing in the objective ℒ. The *W* update is identical to the Lee & Seung (19) KL-divergence update with *H* replaced by *HS*; since *S* is fixed, this is simply the standard update applied to the factorization *V* ≈ *W* (*HS*), and the monotonic decrease guarantee of Lee & Seung applies directly.

For the *H* update, the argument requires a minor extension. Following Lee & Seung, we construct an auxiliary function by bounding log(*W HS*)_*ij*_ from below using Jensen’s inequality. Since (*WHS*)_*ij*_ = ∑ _*a,l*_ *W*_*ia*_*H*_*al*_*S*_*lj*_, we define auxiliary weights over the joint index (*a, l*):

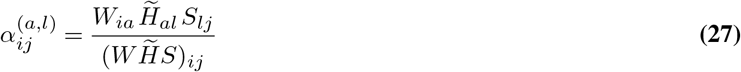

where 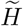 denotes the current iterate. These weights are non-negative and sum to one over (*a, l*) for each (*i, j*), so by the concavity of the logarithm:

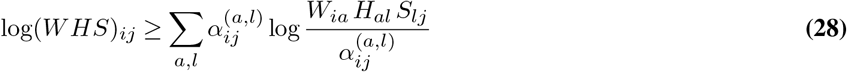

This bound is tight when 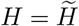. Substituting into ℒ yields an auxiliary function 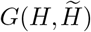 that satisfies *G*(*H, H*) = ℒ (*H*) and 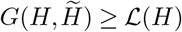. Setting ∂*G/*∂*H*_*al*_ = 0 and solving recovers the multiplicative update in Equation Eq. (21). Since each update minimizes the auxiliary function, ℒ is non-increasing at every iteration:

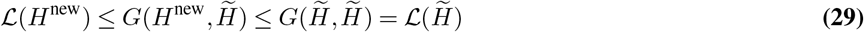

(where H 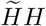 *Hdenotesthecurrentiterate*)”

In practice, convergence is declared when the relative change in ℒ between successive iterations falls below 10^−4^, or when the maximum number of iterations is reached.

## Supplementary Note 3: Description of Competing Methods

To facilitate comparison across methods, we adopt a unified notation. Let *N* denote the number of spatial spots and *G* the number of genes. The observed spatial transcriptomics data are represented by a count matrix 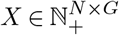, where *X*_*sg*_ denotes the observed expression count of gene *g* in spot *s*. We assume *K* underlying cell types. The proportion of cell type *k* in spot *s* is denoted by *π*_*sk*_, with ***π***_*s*_ = (*π*_*s*1_, …, *π*_*sK*_) lying in the (*K* − 1)-dimensional simplex Δ^*K*−1^. The gene expression profile of cell type *k* is denoted by ***β***_*k*_ = (*β*_*k*1_, …, *β*_*kG*_) ∈ Δ^*G*−1^. Latent discrete cell-type assignments for individual molecules are denoted by *z*_*sm*_. For methods introducing additional latent variables, we retain their original notation but map it to this framework upon introduction.

*Notation correspondence with main text*. The unified notation used in this section relates to the main text as follows: *X* = *V* ^*T*^ (the count matrix is transposed, with spots as rows here and genes as columns), *K* = *k* (number of cell types), and *π*_*s*_ corresponds to the *s*-th column of the normalized 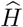 matrix. We adopt the spots-as-rows convention here to match the original publications of the competing methods.

### A. CARD

CARD (7) models spatial correlation in cell-type proportions using a Conditional Autoregressive (CAR) prior. For cell type *k*:

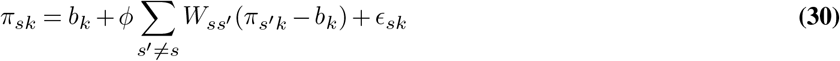

where *b*_*k*_ is the mean proportion of cell type *k* across locations, *W* is a non-negative spatial weight matrix encoding neighborhood structure, *ϕ* controls the strength of spatial autocorrelation, and 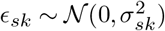. In its standard mode, CARD requires a matched scRNA-seq reference to define cell-type-specific expression profiles and uses a Gaussian likelihood for the spatial transcriptomics data. CARD also provides a reference-free mode that accepts marker gene lists in place of a full single-cell reference; in our benchmark, we used this reference-free mode with marker genes derived from Seurat (see Methods). CARD is therefore included alongside the other methods on equal footing with respect to reference requirements.

### B. STdeconvolve

STdeconvolve (6) applies Latent Dirichlet Allocation (LDA) to spatial transcriptomics data. Each spot is modeled as a mixture of *K* cell types (topics), each characterized by a multinomial distribution over genes. The generative process for spot *s* is:

1. Draw cell-type proportions: *π*_*s*_ ∼ Dirichlet(*α*).
2. For each of the *M*_*s*_ observed molecules in spot *s*:
  a. Draw a cell-type assignment: *z*_*s,m*_ ∼ Multinomial(*π*_*s*_).
  b. Draw a gene: *w*_*s,m*_ ∼ Multinomial 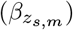.

The posterior distribution over the latent parameters *π* and *β* is:

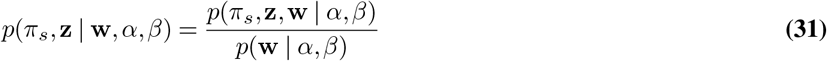

Inference uses variational expectation-maximization. By default, *β* is initialized to zero and *α* = 50*/K*. STdeconvolve preselects genes that are likely to be highly co-expressed within cell types, which can improve deconvolution quality.

### C. SpiceMix

SpiceMix (21) extends NMF with a Hidden Markov Random Field (HMRF) to enforce spatial consistency. Like SNMF, SpiceMix augments the standard NMF framework with spatial regularization; however, SpiceMix learns a spatial precision matrix 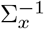 during inference, whereas SNMF uses a fixed spatial mixing matrix **S** computed before optimization.

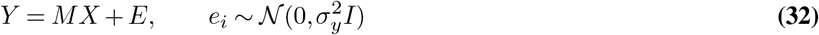

where *M* ∈ ℝ^*G×K*^ contains metagene profiles (columns constrained to the simplex 𝕊_*G*−1_) and 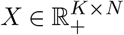 contains per-cell metagene weights. Spatial relationships are represented as a graph 𝒢 = (𝒱, ℰ). The joint likelihood factorizes as:

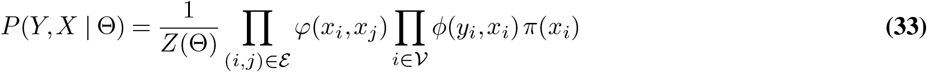

where *ϕ* captures the NMF reconstruction error and *φ* measures spatial affinity between neighboring metagene weight vectors via a learned precision matrix 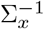:

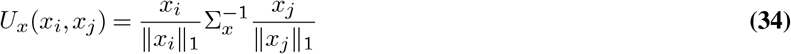

Inference proceeds by coordinate ascent, alternating between estimating the metagene weights *X* (a quadratic program solved via the iterated conditional modes algorithm) and the model parameters Θ (with 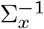 optimized by Adam). The partition function is approximated by Taylor expansion. A regularization hyperparameter *λ*_Σ_ controls the strength of spatial smoothing; as *λ*_Σ_ → 0 the model reduces to standard NMF.

### D. Starfysh

Starfysh (22) is a deep generative model that combines variational inference with a signature-informed Dirichlet prior. Each cell state *k* is represented by a latent centroid *µ*_*k*_ ∈ ℝ^*D*^ (default *D* = 10) and a heterogeneity parameter *α*_*k*_ *>* 0. Each spot *s* is associated with cell-state proportions *π*_*s*_ ∈ Δ^*K*−1^, and its latent representation is:

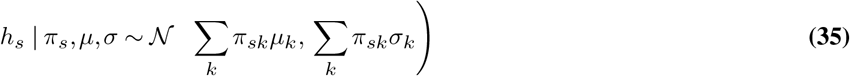

The observed counts are generated from a negative binomial distribution via a neural network decoder *f*:

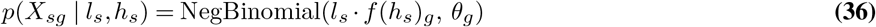

where *l*_*s*_ is the library size and *θ*_*g*_ is a gene-specific dispersion. The prior on *π*_*s*_ is informed by signature scores *A*(*X*_*s*_, *T*_*k*_) computed from user-provided or archetypal marker gene sets:

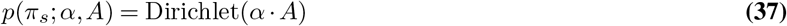

The full generative model is:

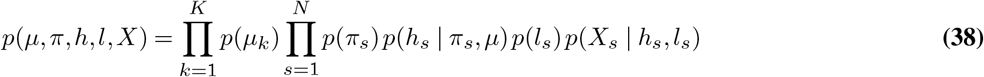

Inference maximizes the ELBO using structured amortized variational inference, with a mean-field approximation over *µ*_*k*_ and amortized encoders for *π*_*s*_ and *l*_*s*_. When marker genes are unavailable, Starfysh uses archetypal analysis to identify the purest spots and derive signature gene sets automatically.

### E. BayesTME

BayesTME (23) models spatial variation in ST data through a hierarchical probabilistic model that simultaneously accounts for cell-type composition, gene expression, spatial smoothing, and spot-level bleeding. The observed count for gene *g* at spot *s* is decomposed across *K* cell types:

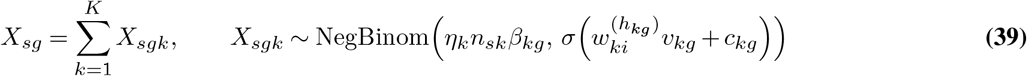

Cell counts per spot are estimated using a graph-fused binomial tree model (GFBT) that enforces spatial smoothness via a first-order graph trend filtering prior on the cell-type probability surface Ψ. The spatial prior is parameterized through the graph Laplacian Δ^(1)^ = Δ^*T*^ Δ, where Δ is the edge-oriented adjacency matrix. Sparsity-inducing horseshoe priors are placed on the gene expression weights. Because full MCMC is computationally intractable for this model, BayesTME uses an empirical Bayes approach that alternates between point estimation of nuisance parameters and full Bayesian inference for the latent variables of interest.

### F. SMART

SMART (24) builds on keyword-assisted topic models (*keyATM*), a class of semi-supervised generative models that integrate prior knowledge in the form of marker genes (keywords) to guide topic formation. In the deconvolution context, spots correspond to documents, genes to words, and cell types to topics. A small set of marker genes per cell type is used to constrain the inference of cell-type proportions and cell-type-specific gene frequencies.

*KeyATM* is based on a mixture of two Dirichlet distributions — one over marker genes only and one over all genes — with the key assumption that marker genes are more highly expressed within their corresponding cell type. The generative process is as follows. Suppose there are *K* cell types, of which the first 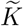 are provided with marker genes.

For each spot *s* and each mRNA molecule *i*:

1. Draw the cell-type assignment: *z*_*si*_ ∼ Categorical(*π*_*s*_).
2. If cell type 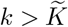 (no marker genes), draw the gene from the standard distribution: *w*_*si*_|*z*_*si*_ = *k* ∼ Categorical(*β*_*k*_), where *β*_*k*_ ∼ Dirichlet(*η*).
3. If cell type 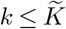 (has marker genes), draw a Bernoulli indicator *d*_*si*_|*z*_*si*_ = *k* ∼ Bernoulli(*ρ*_*k*_), with *ρ*_*k*_ ∼ Beta(*γ*_1_, *γ*_2_). Then:

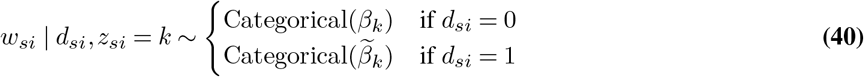

where 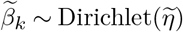 is the marker-gene-specific distribution.
4. The spot-level cell-type proportions follow *π*_*s*_ ∼ Dirichlet(*α*), with

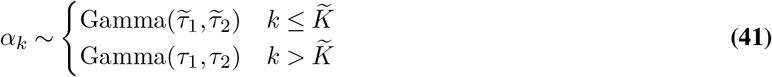

Inference proceeds via collapsed Gibbs sampling with an inverse gene-frequency weighting strategy to prevent highly expressed genes from dominating the inferred topics. The final cell-type-specific gene frequency combines both distributions:

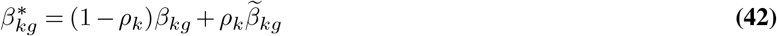

### G. RETROFIT

RETROFIT (25) models the observed count *X*_*sg*_ as a sum of *L* latent components, each decomposed into a background term and a gene-specific term:

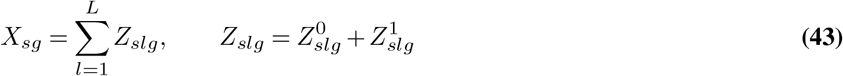

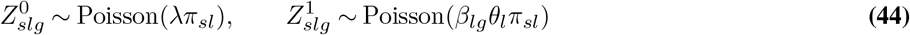

where *β*_*lg*_ is the gene-specific expression level in component *l, θ*_*l*_ is the component contribution, *π*_*sl*_ is the component weight at spot *s*, and *λ* captures a background expression offset. This yields:

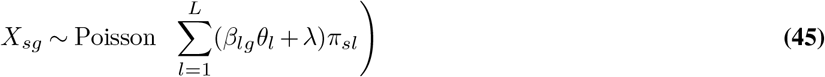

Independent Gamma priors are placed on *β*_*lg*_, *θ*_*l*_, and *π*_*sl*_. Inference uses Structured Stochastic Variational Inference (SSVI), which improves on mean-field variational inference by restoring conditional dependence between model parameters and latent variables. *L* is set to twice the number of known cell types. After inference, components are annotated using provided marker gene sets. In our benchmark, annotation was performed using the same Seurat-derived marker genes provided to other methods that require them (see Methods).

## Supplementary Note 4: Ablation study: implementation details

### Loss Function Variants

Beyond the Kullback-Leibler divergence between two Poisson distributions, we also inspect how well minimizing the Frobenius loss between the observed and reconstructed values *V*_*ij*_ and (*WHS*)_*ij*_ performs. In the latter alternative, the function to minimize is:

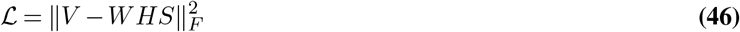

We compute the partial derivative of ℒ with respect to both *H* and *W*:

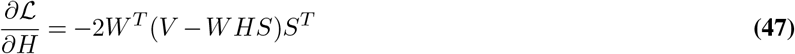

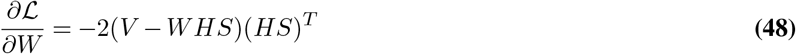

Splitting the positive and negative parts of this gradient, we obtain the following multiplicative update rules:

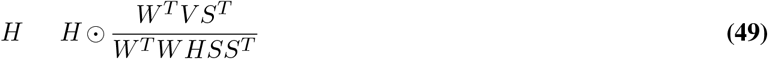

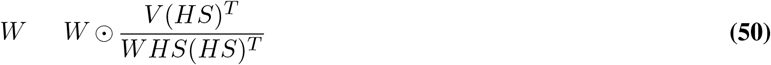

## Supplementary Note 5: Supplementary Figures

**Fig. 4.**
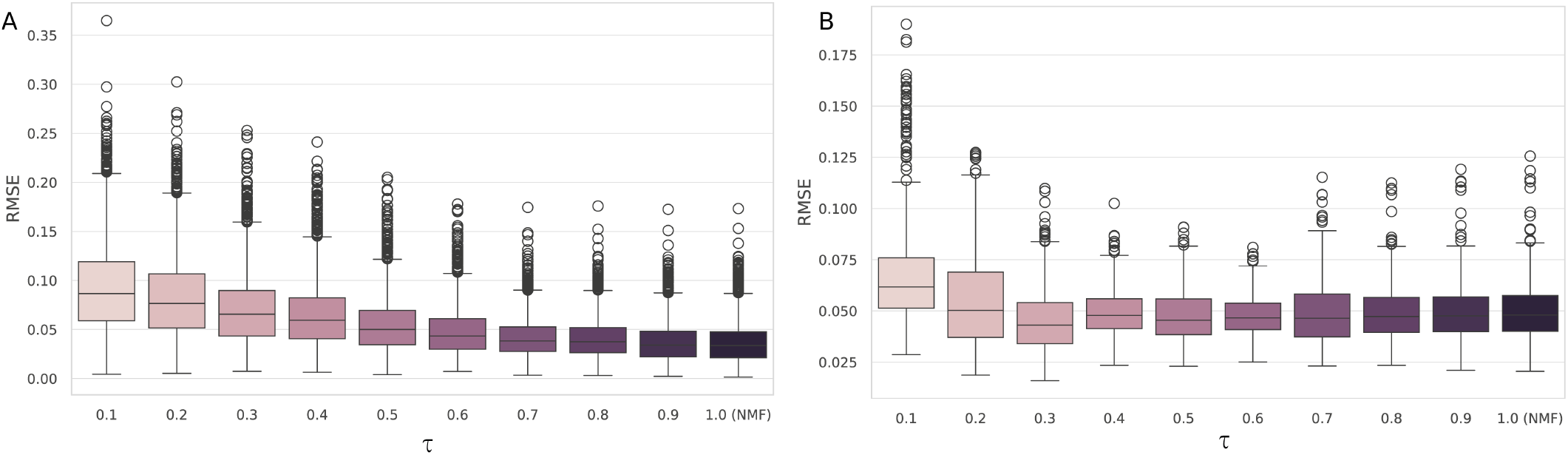
Ablation study: effect of the spatial mixing matrix S on deconvolution accuracy. Root Mean Squared Error (RMSE) distributions across spots are shown for ten values of the mean diagonal of **S**, ranging from 0.1 (strong spatial smoothing) to 1.0 (no smoothing; **S** = **I**, equivalent to standard NMF). **(A)** Results on the TNBC dataset (*k* = 5 cell types). Performance is largely stable across values of the diagonal, and the identity configuration (**S** = **I**) is competitive, suggesting that for small *k* the multiple random restarts employed by SNMF are sufficient to find a good solution without spatial guidance. **(B)** Results on the PDAC dataset (*k* = 20 cell types). Configurations with mean diagonal below 1.0 consistently outperform the identity, with the value of 0.3 yielding the best RMSE. This demonstrates that the spatial mixing matrix acts as a regularizer that is most valuable in high-dimensional factorizations where the search space is large. Box whiskers extend to 1.5*×* the interquartile range.

**Fig. 5.**
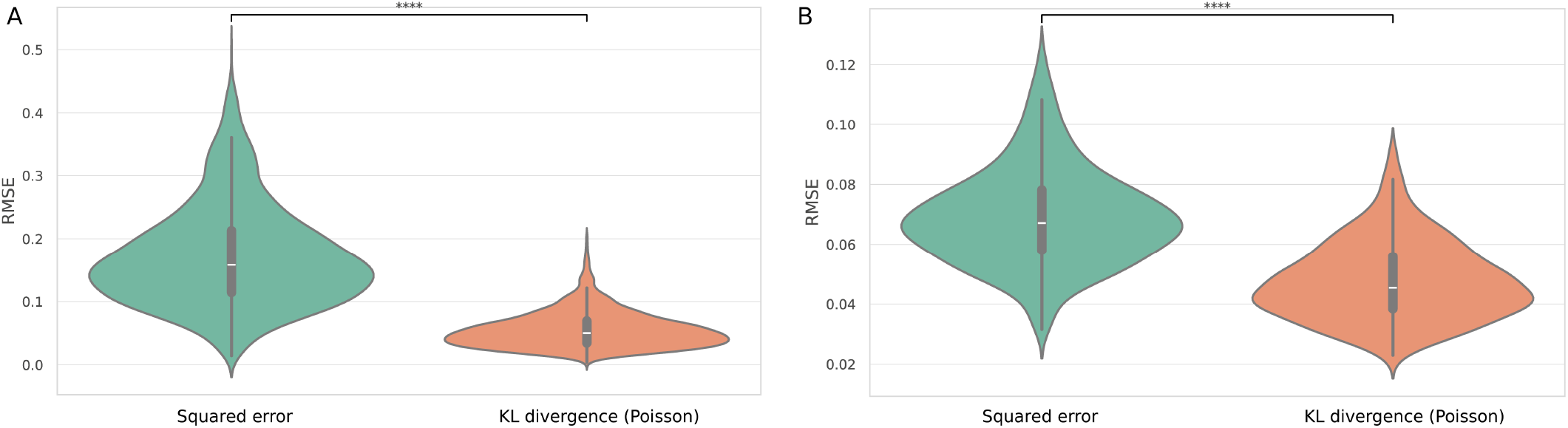
Ablation study: effect of the loss function to optimize ℒ on deconvolution accuracy. Root Mean Squared Error (RMSE) distributions across spots are shown for the Frobenius loss minimization (derivation in Supplementary Section 4), and the Kullback-Leibler divergence minimization (default setting for SNMF). **(A)** Results on the TNBC dataset (*k* = 5 cell types). **(B)** Results on the PDAC dataset (*k* = 20 cell types).

**Fig. 6.**
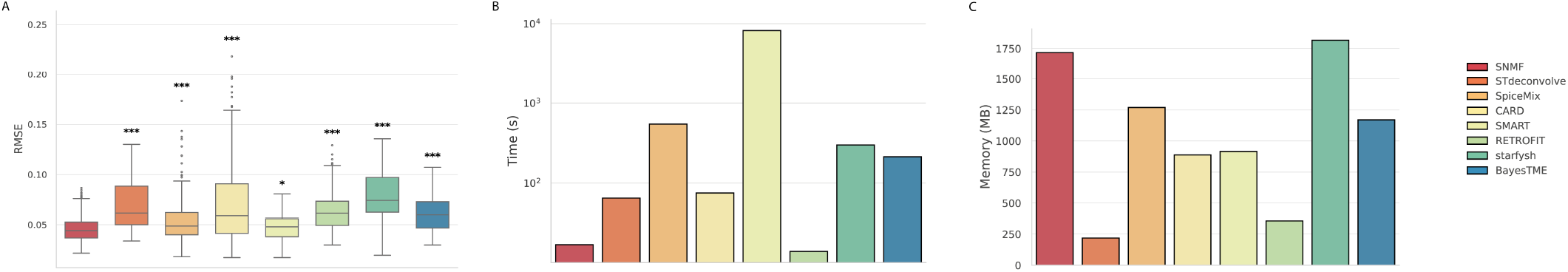
Benchmarking results on the PDAC synthetic dataset (20 cell types). **(A)** Deconvolution accuracy measured by Root Mean Square Error (RMSE) between predicted and ground-truth cell-type proportions across all spots. Each box shows the distribution of per-spot RMSE values for one method. SNMF achieves the lowest median RMSE. Significance levels relative to SNMF are indicated above each box (ns: *p* ≥ 0.05; *: *p <* 0.05; **: *p <* 0.01; ***: *p <* 0.001; Wilcoxon signed-rank test, Bonferroni-corrected). **(B)** Computational runtime (seconds, log scale) for each method on the PDAC dataset, run on the same hardware as the main benchmarking experiments (RTX 3090 Ti with 24 GB RAM). SNMF timings include all 10 random restarts. **(C)** Peak RAM usage (GB) for each method during execution on the PDAC dataset.

**Fig. 7.**
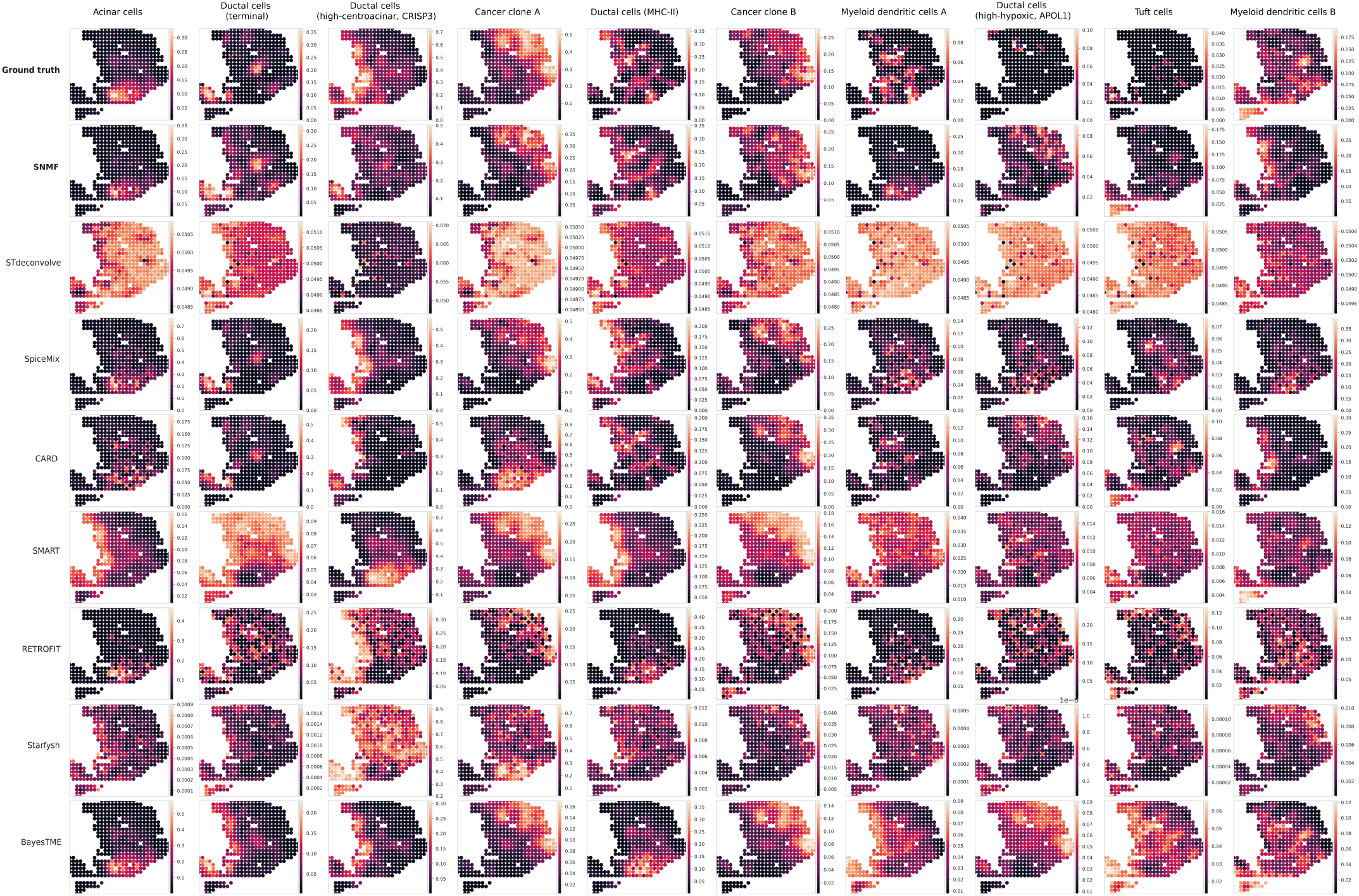
Spatial proportion maps for the PDAC synthetic dataset — cell types 1–10. Each panel shows the predicted proportion of one cell type across all spots for each benchmarked method (columns) alongside the ground truth (first column). Color intensity encodes predicted proportion on the same scale within each row. Results for cell types 11–20 are shown in Supplementary Figure 8.

**Fig. 8.**
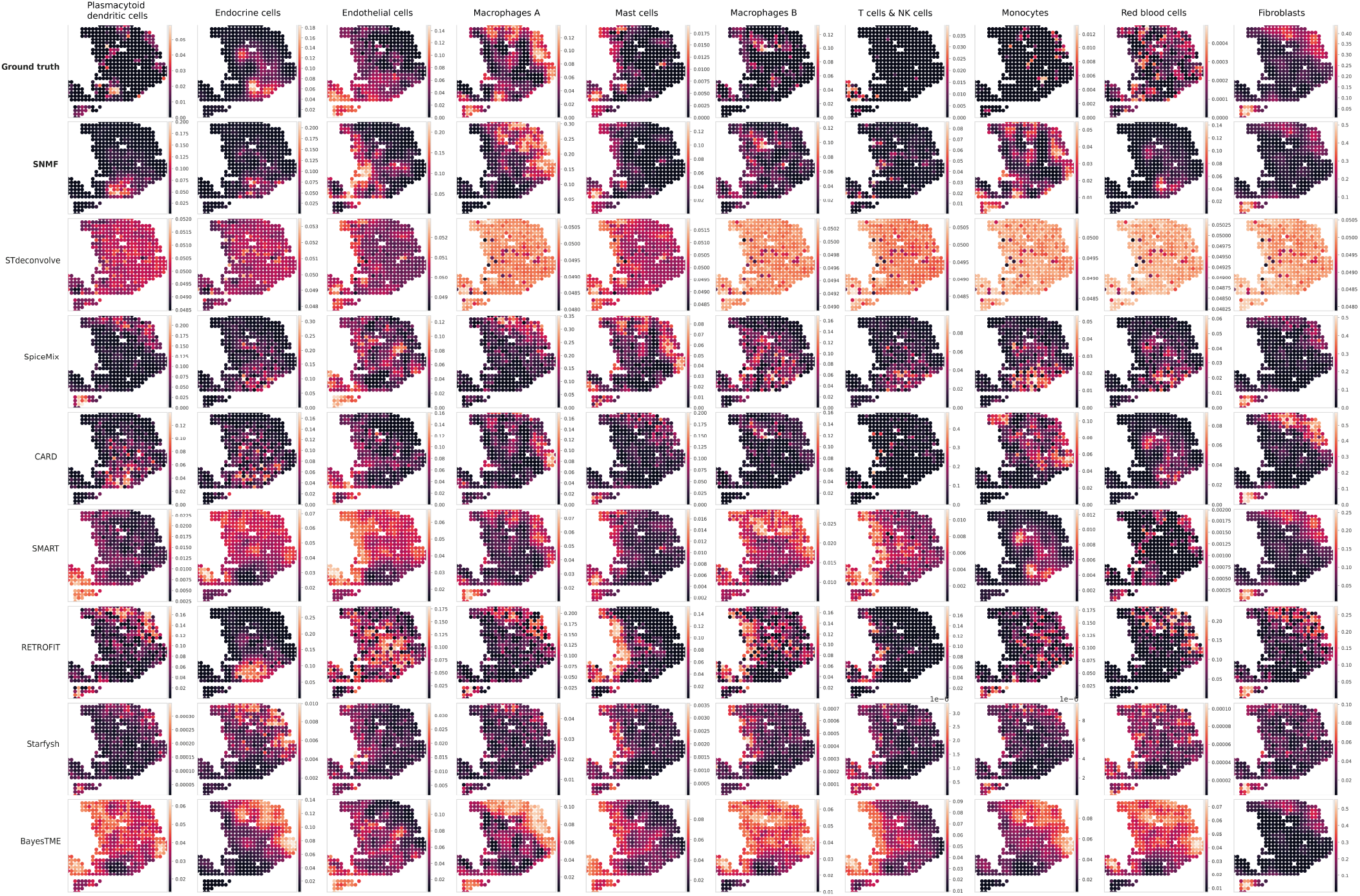
Spatial proportion maps for the PDAC synthetic dataset — cell types 11–20. Layout as in Supplementary Figure 7.

**Fig. 9.**
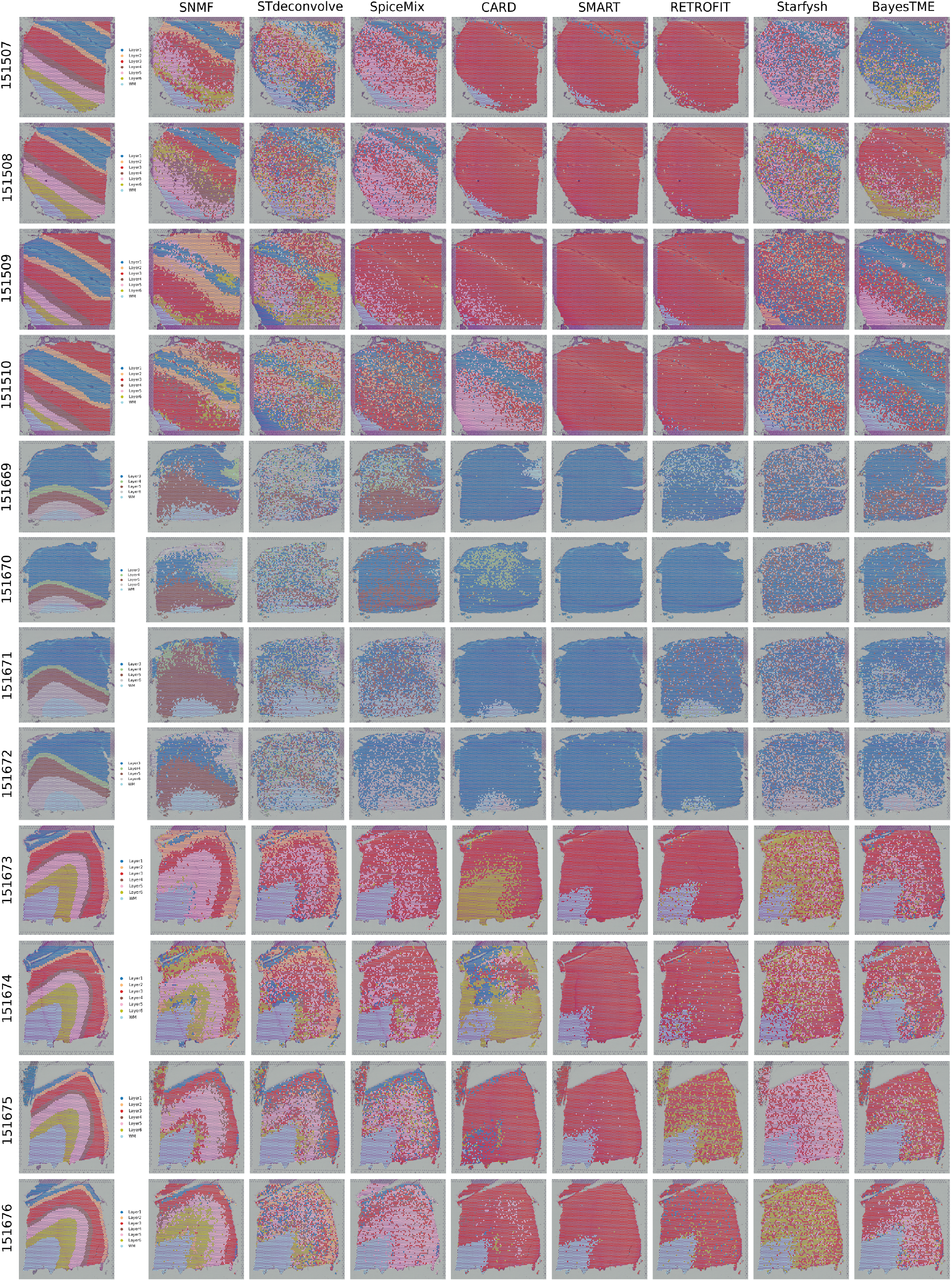
Spatial maps of predicted and annotated dominant cell types across all 12 DLPFC tissue sections. For each section, spots are colored by their dominant cell type as predicted by SNMF (top row) and as assigned by the expert manual annotation (bottom row). Cortical layers (L1–L6) and white matter (WM) are distinguished by color. Sections 151669–151672 lack Layer 1 annotation and were analyzed with *k* = 5; all other sections were analyzed with *k* = 7. Sections are ordered by sample ID (151507–151676). The ARI for each section is shown below each panel. SNMF best recovers the laminar organization of the cortex in 11 out of 12 sections; In section 151671 SNMF ranks second.

## Supplementary Note 6: Supplementary Tables

**Table 2.**
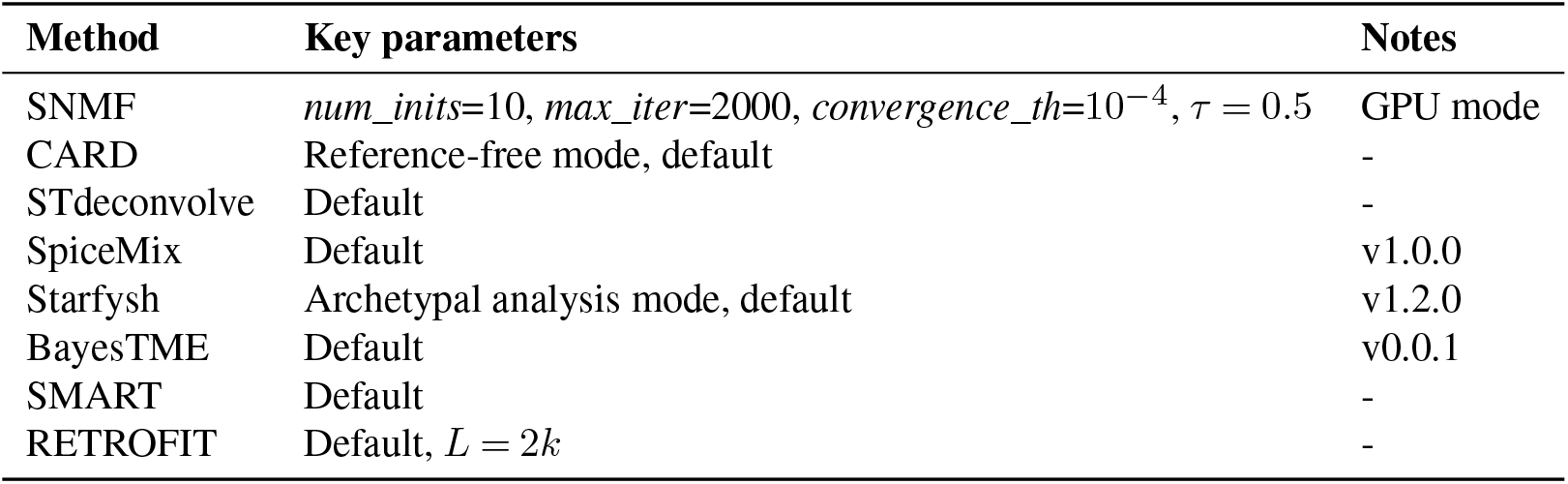
Parameters used for each benchmarked method. All methods were run with default parameters unless otherwise noted. The number of cell types *k* was set to the ground-truth value for each dataset (see Supplementary Table 4 for per-section DLPFC values).

**Table 3.** Marker genes used for methods requiring prior gene information. Marker genes were identified using Seurat’s `FindAllMarkers` function (36) with parameters (e.g., *logfc*.*threshold* = 0.25, *min*.*pct = 0*.*25, test*.*use = “wilcox”*). The statistically siginficant marker genes per cell type were retained. The same marker gene sets were provided to all methods that accept them (CARD in reference-free mode, STdeconvolve, Starfysh, SMART and RETROFIT). For the TNBC and PDAC synthetic datasets, marker genes were derived from the scRNA-seq references used to generate the synthetic data. For the DLPFC dataset, marker genes were derived from the spatial data itself. The full marker gene lists are provided in Zenodo (https://doi.org/10.5281/zenodo.18852117).

| Dataset | Sample | Cell types | Avg. number of markers per cell type |
| --- | --- | --- | --- |
| TNBC | - | 5 | 30.2 |
| PDAC | - | 20 | 378.35 |
| DLPFC | 151507 | 7 | 508.42 |
| DLPFC | 151508 | 7 | 546.42 |
| DLPFC | 151509 | 7 | 777.85 |
| DLPFC | 151510 | 7 | 618.14 |
| DLPFC | 151669 | 5 | 1,522.00 |
| DLPFC | 151670 | 5 | 1,247.40 |
| DLPFC | 151671 | 5 | 1,243.60 |
| DLPFC | 151672 | 5 | 891.80 |
| DLPFC | 151673 | 7 | 919.75 |
| DLPFC | 151674 | 7 | 996.00 |
| DLPFC | 151675 | 7 | 765.00 |
| DLPFC | 151676 | 7 | 552.14 |

**Table 4.** Summary of the 12 DLPFC tissue section datasets. Genes were filtered to retain only those expressed in at least 10% of spots within each section. The number of annotated regions reflects the expert-annotated cortical layers and white matter present in each section; sections 151669–151672 lack Layer 1 annotation and therefore have 5 regions rather than 7. The value of *k* used for all methods matches the number of annotated regions in each section.

| Sample ID | Spots | Genes after filtering | Annotated regions ($k$) |
| --- | --- | --- | --- |
| 151507 | 4,226 | 3,975 | 7 |
| 151508 | 4,384 | 3,220 | 7 |
| 151509 | 4,789 | 4,130 | 7 |
| 151510 | 4,634 | 3,856 | 7 |
| 151669 | 3,661 | 5,216 | 5 |
| 151670 | 3,498 | 4,866 | 5 |
| 151671 | 4,110 | 5,428 | 5 |
| 151672 | 4,015 | 5,084 | 5 |
| 151673 | 3,639 | 6,542 | 7 |
| 151674 | 3,673 | 7,798 | 7 |
| 151675 | 3,592 | 5,438 | 7 |
| 151676 | 3,460 | 5,780 | 7 |

**Table 5.** Adjusted Rand Index (ARI) for all methods across all 12 DLPFC tissue sections. The best-performing method for each sample is shown in bold. SNMF achieves the highest ARI in 11 out of 12 samples. In sample 151671, where SNMF does not rank first, it ranks second. Mean and standard deviation across all 12 samples are reported in the final two rows. Absolute ARI values are moderate for all methods, reflecting the inherent difficulty of evaluating deconvolution via dominant cell-type assignment in a tissue where each annotated region contains a mixture of cell types (see main text for discussion). Sections 151669–151672 were analyzed with *k* = 5 (lacking Layer 1 and 2 annotation); all others with *k* = 7.

| Sample | SNMF | STdeconvolve | SpiceMix | CARD | SMART | RETROFIT | Starfysh | BayesTME |
| --- | --- | --- | --- | --- | --- | --- | --- | --- |
| 151507 | **0.382** | 0.087 | 0.143 | 0.032 | 0.015 | 0.007 | 0.033 | 0.117 |
| 151508 | **0.324** | 0.074 | 0.070 | 0.040 | 0.000 | 0.004 | 0.021 | 0.133 |
| 151509 | **0.285** | 0.088 | 0.119 | 0.075 | 0.000 | 0.039 | 0.023 | 0.211 |
| 151510 | **0.244** | 0.060 | 0.070 | 0.162 | 0.000 | 0.008 | 0.045 | 0.149 |
| 151669 | **0.246** | 0.047 | -0.059 | -0.051 | 0.000 | -0.085 | -0.001 | 0.204 |
| 151670 | **0.169** | 0.012 | 0.083 | -0.078 | 0.000 | -0.003 | -0.018 | 0.162 |
| 151671 | 0.136 | 0.095 | 0.022 | 0.091 | 0.000 | 0.088 | 0.066 | **0.212** |
| 151672 | **0.335** | 0.042 | 0.080 | 0.079 | -0.005 | 0.044 | 0.055 | 0.096 |
| 151673 | **0.430** | 0.230 | 0.150 | 0.203 | 0.098 | 0.112 | 0.073 | 0.190 |
| 151674 | **0.313** | 0.277 | 0.152 | 0.103 | 0.070 | 0.036 | 0.076 | 0.197 |
| 151675 | **0.351** | 0.331 | 0.216 | 0.021 | 0.088 | 0.141 | 0.089 | 0.190 |
| 151676 | **0.363** | 0.203 | 0.144 | 0.086 | 0.104 | 0.133 | 0.106 | 0.188 |
| Mean | **0.298** | 0.129 | 0.099 | 0.064 | 0.034 | 0.044 | 0.047 | 0.171 |
| Std | 0.083 | 0.099 | 0.069 | 0.076 | 0.044 | 0.062 | 0.036 | 0.037 |

**Table 6.**
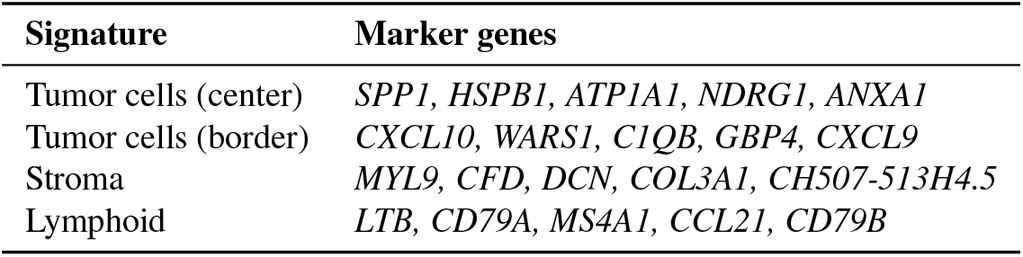
Gene signatures used to annotate the cell-type components inferred by SNMF for the melanoma dataset, obtained from Zhao et al. (35).

## Notes

https://github.com/ML4BM-Lab/SNMF-paper/

https://doi.org/10.5281/zenodo.18852117

### Competing Interest Statement

The authors have declared no competing interest.

